# The selective dynamics of interruptions at short tandem repeats

**DOI:** 10.1101/2025.06.09.658724

**Authors:** Michael E. Goldberg, Harriet Dashnow, Kelley Harris, Aaron R. Quinlan

## Abstract

Short tandem repeats (STRs) are hotspots of genomic instability that mutate at rates orders of magnitude greater than non-repetitive loci due to frequent replication slippage. Expansions at some STR loci cause Mendelian diseases, while variation at other noncoding loci may affect complex traits, possibly by altering transcription factor occupancy of nearby binding sites. Accordingly, some STRs are inferred to be under purifying selection, regardless of their instability. One or more “interruptions”, or bases that disrupt the locus’s canonical repeat, significantly decrease an STR’s mutability. For example, the onset of Huntington’s Disease, a neurodegenerative disorder associated with somatic expansions of a trinucleotide coding STR, is delayed in individuals whose inherited alleles contain interruptions. Thus, interruptions that decrease mutation rate at some coding loci may broadly protect against deleterious phenotypes associated with locus instability. However, interruptions may themselves be deleterious at constrained loci, particularly at noncoding loci in gene regulatory elements, possibly disrupting the formation of secondary structures key to their function. We therefore hypothesized that the frequency of interruptions could depend on a locus’s functional importance–at constrained loci, the fitness effects of expansions but also interruptions could be more deleterious than at neutral loci. To test this hypothesis, we examined the distribution of interruptions at ∼650,000 autosomal STRs. In the ∼2,500 3- or 6bp-motif coding STRs, we find that synonymous interruption density increases with purifying selection on the gene, while the opposite is true for missense-causing interruptions. In contrast, noncoding STRs in gene regulatory elements harbor fewer interruptions than those unassociated with function and putatively evolving neutrally. Our findings indicate that the abundance of interruptions may be partially explained at coding STRs by the benefit of lower instability. In contrast, maintaining a minimum core stretch of uninterrupted repeat may be key to the function of noncoding STRs that fall within regulatory elements, outweighing the benefits of stability.

## Introduction

Short tandem repeats (STRs) are repetitive elements that comprise tandem 1-6 bp motifs and account for 3% of the human genome (International Human Genome Sequencing Consortium et al. 2001). In the pre-genomic era, STRs were commonly used as population markers and for DNA fingerprinting thanks to their high rates of heterozygosity (Weber and Wong 1993; Spencer et al. 2000). Their high heterozygosity results from remarkable genomic instability; STRs mutate at rates orders of magnitude greater than non-repetitive loci due to frequent replication slippage, most recently estimated around 5.5 * 10^−6^ mutations/site*generation (Weber and Wong 1993; Sun et al. 2012; Willems et al. 2017; Mitra et al. 2021; Steely et al. 2022; Kristmundsdottir et al. 2023; Porubsky et al. 2024). Due to their repetitive nature, STRs are more likely than high complexity genomic sequence to form bulky, non-B conformations of DNA, such as hairpins or cruciforms (Trinh 1991; Khristich and Mirkin 2020; McGinty and Sunyaev 2023 Mar 13). These bulky structures likely contribute to replication fork stalling, which results in slippage of the nascent strand. Slippage may result in contraction or expansion of an allele by one or more motifs relative to the template strand. Though slippage resulting in mutagenesis is typically linked to cell division, STRs may also expand or contract outside of S-phase replication (Goldberg et al. 2024; Handsaker et al. 2025).

A handful of local characteristics are known to determine an STR’s mutation rate. Longer alleles, shorter motifs, and lower GC content are generally associated with higher mutation rate; other characteristics and loci that act in *cis* or *trans* have also been described (Brinkmann et al. 1998; Legendre et al. 2007; Sun et al. 2012; Ananda et al. 2013; Kristmundsdottir et al. 2023; McGinty and Sunyaev 2023 Mar 13). Interruptions, or allelic impurity, refer to substitutions or small insertions or deletions that disrupt an otherwise pure repeat motif and are known to slow down the rate of slippage (Gacy et al. 1995; Sainudiin et al. 2004; Ananda et al. 2014; Wright et al. 2019; McGinty and Sunyaev 2023 Mar 13; Rajan-Babu et al. 2024; Danzi et al. 2025; McGinty et al. 2025). Though the exact mechanism is unclear, interruptions may decrease the likelihood of forming non-B conformations and therefore the expected rate of replication stalling and slippage (Gacy et al. 1995; Pearson et al. 1998). As such, mutation rate roughly scales with the longest remaining stretch of pure repeat. Although single base interruptions may be caused by mutational processes independent of STRs, nucleotides added during STR expansion are erroneous at a rate of 10^−2^ to 10^−3^ per added nucleotide; contractions may also delete interruptions (McGinty and Sunyaev 2023 Mar 13).

Variation at 60 STR loci in coding exons has been long associated with a number of Mendelian diseases typically involving neurodegeneration linked to somatic expansion following inheritance of an allele at or beyond pathogenic length (Hiatt et al. 2025). Huntington’s Disease (HD), for example, is linked to somatic expansions at a CAG trinucleotide repeat encoding a polyglutamine stretch in the *HTT* gene (Wright et al. 2019). Individuals who inherit an allele that contains 40 CAG motifs or greater are at risk of disease; the greater the length, the earlier the age of onset due to the higher slippage rate per unit time. However, individuals who inherit an allele with a CAG>CAA interruption have a significantly increased age of onset conditional on the length of the polyglutamine. Similarly, interspersed AGG repeats stabilize *FMR1* and decrease the probability of intergenerational expansion that leads to Fragile X syndrome (Eichler et al. 1995; Eichler and Nelson 1996). Broadly, interruptions stabilize the repeat, protecting it from the slippage that results in deleterious phenotypes (Chen et al. 2024; Rajan-Babu et al. 2024).

Intronic and intergenic noncoding STRs were historically assumed to be largely nonfunctional, outside of CG-rich repeats at which methylation is associated with gene regulation. However, a number of recent studies have reported noncoding STR variation has causal links with a number of complex traits such as developmental disorders, cancer risk, and height (Gymrek et al. 2016; Fotsing et al. 2019; Mukamel et al. 2021; Mukamel et al. 2023; Grasberger et al. 2024). Variation at some STRs directly causes variance in gene expression (Fotsing et al. 2019; Margoliash et al. 2023; Tanudisastro et al. 2025 Apr 9). A proposed mechanism behind these molecular and trait associations is that STRs may alter gene expression when proximal to the binding motif of transcription factors and other DNA binding proteins (Horton et al. 2023). In vitro work has shown that these proteins can directly bind STRs, thus increasing the binding occupancy of a neighboring motif at an enhancer or other *cis*-acting gene regulatory element. Different proteins bind more strongly and for a longer amount of time to different STR motifs; evolutionary similarity between proteins does not always predict similar motif preferences. Although purity broadly correlates with stronger binding, the effect is not monotonic (Horton et al. 2023). Accordingly, a proportion of these noncoding STRs are inferred to be evolving under purifying selection.

Identifying constraint at STR loci has historically been challenging due to their rapid mutagenesis and high diversity. Many methods that identify constrained high-complexity genomic regions as those that are depleted for SNP variation under a neutral model of their mutagenesis (Lek et al. 2016; Havrilla et al. 2019; Karczewski et al. 2020; Chen et al. 2022). LOEUF, for example, quantifies constraint at transcripts and genes by estimating the depletion of observed loss-of-function alleles relative to neutral expectation (Karczewski et al. 2020). This score represents a locus’s intolerance to dominant loss-of-function. New models that directly identify STR constraint similarly rely on models of their complex mutagenesis and require making similar assumptions of dominance (Gymrek et al. 2017; Mitra et al. 2021; Haasl and Payseur 2024).

The selective dynamics of a genetic element that controls mutation rate were initially derived by Kimura in 1967 (Kimura 1967). A ‘mutator’ is a genetic element that controls the mutation rate of a genomic region or set of elements; excluding pleiotropy beyond its effect on germline mutation rate, a mutator allele’s selection coefficient scales with the change in mutation rate, the distribution of fitness effects of those mutations, and the linkage to the mutations. Interruptions stabilize an STR allele in a way that mirrors a *cis*-acting ‘anti-mutator’ in perfect linkage to the mutations whose rate it decreases. If slippage at a locus is deleterious, interruptions that stabilize an allele may under indirect positive selection. However, interruptions may themselves be deleterious and under direct purifying selection at constrained loci, particularly at noncoding loci in gene regulatory elements, disrupting the purity possibly key to their function. We would hypothesize, therefore, that the density of interruptions at STR loci could scale with constraint. At loci evolving entirely neutrally, expanded and contracted alleles and pure and interrupted alleles are as fit as an unmutated allele, so interruptions that increase stability and decrease purity could accumulate at a rate only affected by drift. However, at loci evolving under constraint, we could see either of two effects: if expanded and contracted alleles are more deleterious than an interrupted allele of the ancestral length, constrained loci could be enriched for interruptions. The opposite could occur if impurity is more deleterious than expanded and contracted alleles. Furthermore, this effect could scale monotonically with constraint and be strongest at loci where purifying selection is highest and intermediate where constraint is mild but nonzero.

To test this hypothesis, we examined the distribution of interruptions at the major alleles of ∼650,000 human autosomal STRs in large population databases. In the ∼2,500 STRs that fall within coding exons and comprise 3- or 6-bp motifs whose expansions and contractions would not change reading frame, we find that the density of interruptions in coding STRs increases with purifying selection on the gene, provided those interruptions lead to no amino acid change. In contrast, noncoding STRs under purifying selection harbor fewer interruptions than those evolving neutrally. Our findings indicate that the abundance of interruptions may be partially explained at coding STRs by the benefit of a lower mutational burden at their linked loci. In contrast, maintaining a minimum core stretch of uninterrupted repeat may be key to the function of noncoding STRs that fall within regulatory elements, outweighing the benefits of lowering the mutation rate.

## Results

### Coding STRs in constrained genes are enriched for silent interruptions

Interruptions that stabilize an STR allele at a small number of disease-related loci are protective against deleterious phenotypes associated with their slippage (Eichler et al. 1995; Wright et al. 2019; Chen et al. 2024). We hypothesized that this phenomenon may generalize genome-wide and that interruptions would be favored at coding loci under constraint. However, at loci evolving neutrally whose alleles are equally fit regardless of their length, we hypothesized that interruptions would not be selectively beneficial and accumulate under a neutral model of mutagenesis. If this hypothesis is correct, we would observe that locus constraint covaries with its density of interruptions.

To test this, we examined 2838 trinucleotide and hexanucleotide STR loci whose allele in the GRCh38 reference genome was fully contained in an exon from a canonical transcript in RefSeq Select or in a medically-relevant or constrained exon in Matched Annotation from the NCBI and EMBL-EBI (MANE) (Morales et al. 2022; Goldfarb et al. 2025). STR genotypes from 2852 individuals of diverse ancestry, comprising all unrelated individuals from the 1000 Genomes (1KGP) and the H3Africa (H3A) cohorts, were previously generated using HipSTR (Jam et al. 2023). These STRs are short enough to be fully spanned by one end of an Illumina short read and thus represent a certain subset. We filtered loci that were missing genotypes at >25% of individuals, failed Hardy-Weinberg Equilibrium (*p* < 0.000001), or overlapped a known segmental duplication and applied sample-level filters as described in (Jam et al. 2023) and identified the major allele amongst those remaining. Coding major alleles spanned lengths of 14 - 104 bp.

To count the number of interruptions for each major allele, we took an evolutionary approach assuming that, at some point in time before the observed major allele was derived, this “ancestral” version of the major allele was pure and that interruptions accumulated over time (Supplemental Figure 1). This method likely overestimates the number of interruptions, but we assume that the degree of overestimation is independent of locus constraint so will not affect our models. We inferred a hypothetical pure allele composed of the locus’s motif as detected by HipSTR and of identical length to the observed major allele that minimized the number of amino acid changes between the two and excluded nonsense mutations over the theoretical evolutionary history (**Methods**). For each codon, we compared the nucleotide sequence between the hypothetical pure ancestral allele and determined whether there were any substitutions (i.e., was the codon interrupted), and whether the substitution(s) resulted in a synonymous or missense change. Codons differ in their synonymous “mutational space”, i.e. of all nine possible individual single nucleotide substitutions, a different number of them lead to silent mutations. To account for these differences in mutational space, we calculated the ratio of all possible synonymous to nonsynonymous single base substitutions from the ancestral allele. Alleles composed of 25% interrupted bases were excluded. We used the containing gene’s LOEUF score as a proxy for locus constraint; LOEUF represents a gene’s intolerance to dominant loss-of-function mutation. Low LOEUF values correspond to strong purifying selection, signaled by a depletion of segregating deleterious variants.

**Supplemental Figure 1:**
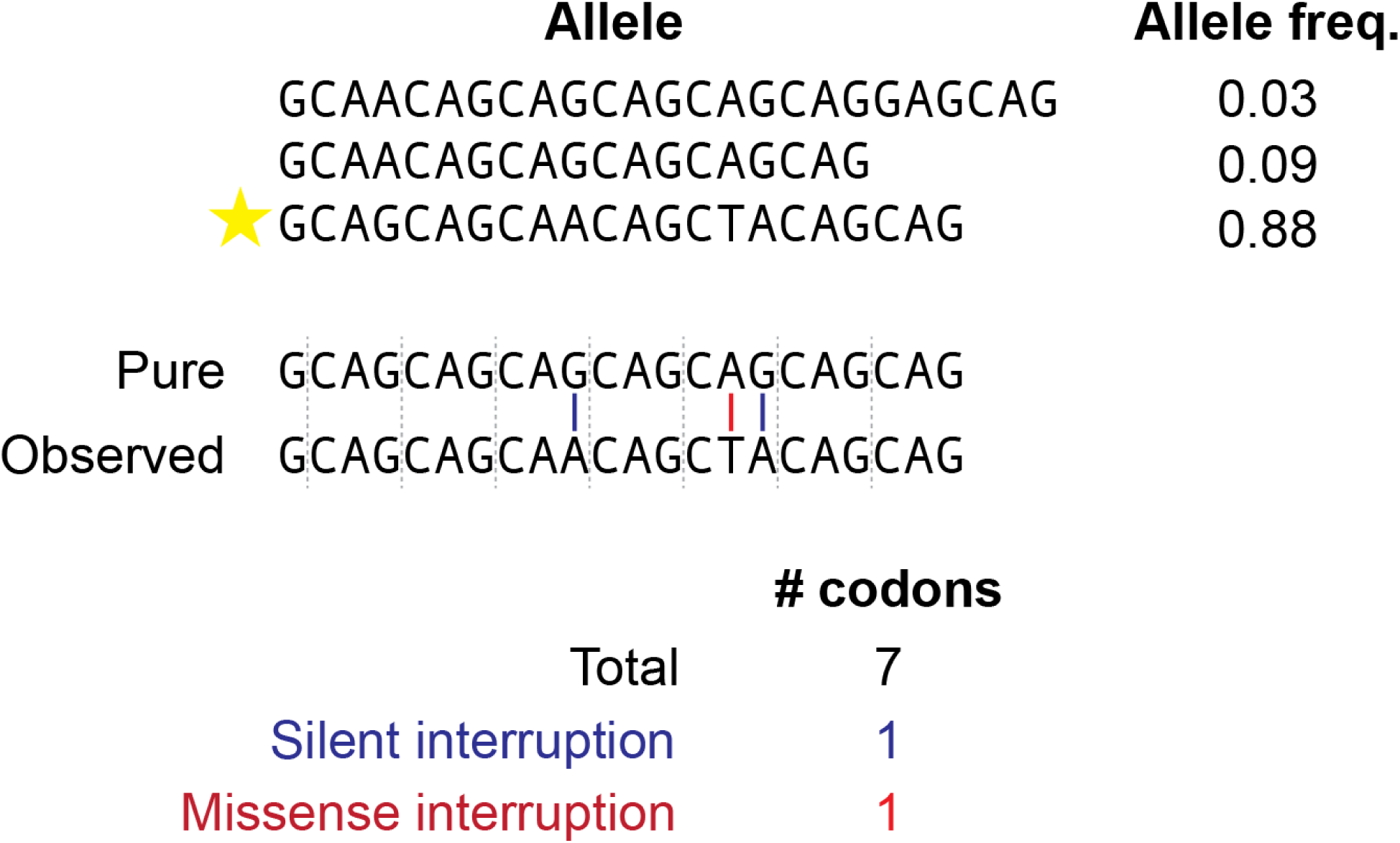
Schema for determining the number of interruptions at coding STRs at a hypothetical polyglutamine repeat. At each locus, we selected the major allele and determined the number of substitutions between the observed allele and a pure repeat of the same length. We classified each codon as being pure or interrupted. Silently interrupted codons only contained synonymous substitutions, while any missense substitution classified a codon as missense. Alleles with nonsense substitutions were excluded from the dataset.

Of the 2554 STR loci that fall within genes and are annotated with a LOEUF score, the majority of major alleles are interrupted relative to a hypothetical pure allele (66.1%). LOUEF scores of the containing genes range from 0.051 - 1.978. To determine the effects of constraint on interruptions, we built a series of Poisson generalized linear models (GLMs) with log link functions. These models included two covariates to control for characteristics that are known to affect locus stability: motif length and GC content. We controlled for allele length and mutational space as an offset variable. After correcting for these characteristics, we observed that major alleles are enriched for silent over missense interruptions (Poisson GLM with log link, *p* < 2.2 * 10^−16^; **Figure 1A**; see **Methods**). Unless otherwise specified, all GLMs in this manuscript correct for motif length and GC content as covariates. In this section, mutational space and allele length are included in an offset variable.

**Figure 1.**
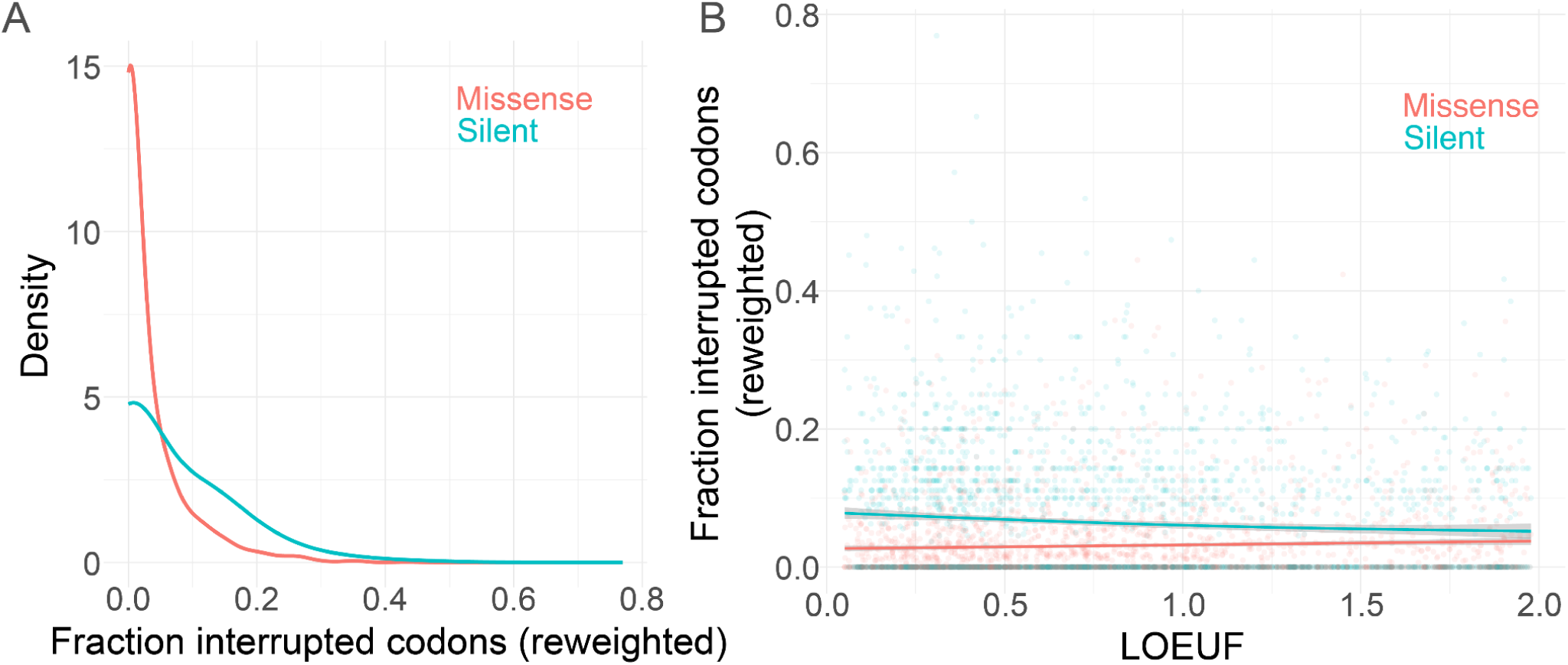
Interruptions at coding STRs. **A**: Silent interruptions are common and missense interruptions are rare at constrained coding STRs. The numbers of missense and silent interruptions are reweighted based on mutational space. **B**: Silent interruptions are enriched at constrained genes while missense interruptions are depleted. The fractions of interrupted codons are reweighted by mutational space of silent and missense interruptions, respectively. Fraction of interrupted codons for the major allele at each coding STR is plotted as a function of the containing gene’s LOEUF score. Lower LOEUF scores correspond to genes less tolerant of dominant loss-of-function alleles, thus indicating stronger purifying selection.

The number and consequence of interruptions covary significantly with genic constraint. STRs within highly constrained genes have significantly more silent interruptions than STRs in genes evolving more neutrally (Poisson GLM, *B* = -0.24, *p* = 5.34 * 10^−8^). In contrast, constraint nominally decreases the number of missense-causing interruptions (Poisson GLM, *B* = 0.071, *p* = 0.053). A GLM with an interaction term finds that silent interruptions are significantly more prevalent than missense interruptions and that constraint has a significantly different effect on the number of interruptions conditional on their coding consequence (Poisson GLM; *B =* 1.40, -0.41; *p* < 2 * 10^−16^, *p* = 1.14 *10^−12^, respectively; **Figure 1B**). Modelling constraint significantly improves a Poisson GLM, indicating that constraint helps explain coding STR purity (ΔAIC = -45).

We examined how alternate measures of coding STR constraint covaried with interruptions and found heterogeneous effects. To independently approximate constraint, we examined whether falling in a known autosomal dominant (AD) gene affected interruption prevalence (Berg et al. 2013). The 159 loci that fall in AD genes are not significantly differently interrupted than the 2644 loci that do not fall in these genes, regardless of the protein coding consequence of the interruption (Poisson GLM, *p* = 0.478). Relatively few loci fell in AD genes, potentially limiting our power to detect any effect on allele purity. We next tested whether an STR locus overlapping an annotated protein domain affected its purity. Given our previous findings described in Figure 1, we expected to see that STRs in protein domains contained more silent and fewer missense interruptions than STRs outside of annotated domains. Contrary to these expectations, however, the 317 STR loci that fall within known protein domains have significantly fewer silent interruptions (Poisson GLM, *p* = 1.14 * 10^−6^). The effect of genic constraint on silent interruptions does not significantly differ between STRs within and outside of protein domains (Poisson GLM, *p* = 0.13).

Broadly, we found evidence that selective constraint affects purity at trimer and hexamer STRs contained in genes. STRs in genes under strong constraint are significantly more likely to harbor interruptions, provided that those interruptions arise from mutations that did not change the protein coding sequence. On the other hand, missense-causing interruptions were less common at constrained genic STRs. This interaction supports a hypothesis that interruptions may be selectively beneficial at highly constrained loci by decreasing the local rate of deleterious expansion and contraction through slippage. However, the direct purifying selection of a missense variant may outweigh the indirect positive selection granted by slowing slippage rates, leading to significantly different associations between interruptions and constraint while considering the protein coding consequence.

### Constraint and functional heterogeneously affect purity at noncoding STRs

We then asked if purity was similarly affected by constraint at noncoding STRs. Most noncoding STRs are likely evolving neutrally or nearly neutrally in humans, but a subset is hypothesized to be under constraint, likely thanks to their capacity to affect gene regulation (Gymrek et al. 2017; Mitra et al. 2021; Horton et al. 2023). STRs can modify the affinity of DNA binding proteins when they flank the protein’s binding motif; the neighboring STR is directly bound by the protein before binding to the motif, thus increasing the amount of time the motif is bound. STRs are further known to be enriched in enhancers (Sawaya et al. 2013; Gymrek et al. 2016; Horton et al. 2023).

Expansion and contraction appear to be deleterious at these noncoding loci, so we hypothesized that slowing the causal slippage by decreasing locus purity may be selectively beneficial, as for coding STRs (Mitra et al. 2021). If purifying selection on expanded or contracted alleles is strong, we could expect to see an enrichment for interruptions at constrained noncoding STRs. However, interrupting an STR that flanks a binding motif can heterogeneously decrease the motif’s binding occupancy (Horton et al. 2023). Thus, the direct functional consequence of a substitution could possibly outweigh the benefits of slowing slippage. If purity is more selectively beneficial than instability is deleterious, constrained noncoding STRs could be depleted for interruptions.

Similar to the analyses above, we considered the purity of the major allele only at 644,070 nonhomopolymer noncoding STRs. Interruptions were quantified slightly differently than at coding repeats: unable to consider the consequence of each interruption on an amino acid sequence, we simply calculated the minimum Levenshtein distance to a hypothetical pure ancestral version of the repeat composed entirely of the reference repeat unit. We used a modified alignment approach to account for indels in inferring an ancestral pure allele (**Methods**). 61.3% of loci harbored at least one interruption (**Supplemental figure 2**). As before, we excluded loci whose major alleles were more than 25% interrupted (Levenshtein distance / allele length in bp) as these alleles could be incorrectly genotyped or include multiple major motifs; this roughly corresponded to the 99^th^ percentile of impurity.

**Supplemental Figure 2:**
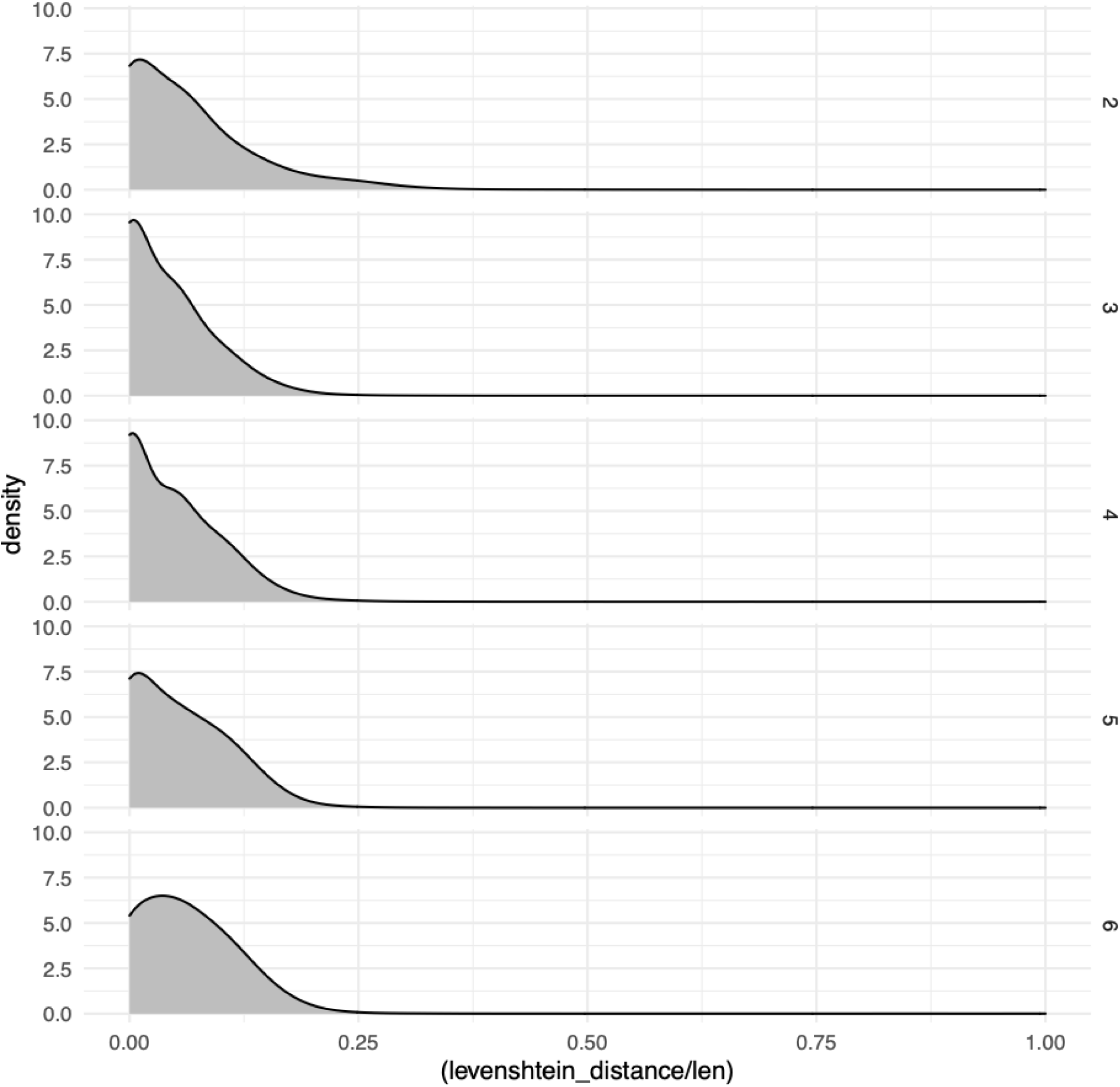
Noncoding STR purity by repeat unit length.

Both STR constraint and the functional consequence of an interruption are trickier to quantify at noncoding loci so we relied on several different annotations. We first examined interruptions across the noncoding genome as a function of their chromatin structure and expected function in gene regulation as identified by chromHMM. To best approximate the chromatin state in the germline, we used 25-state chromHMM annotations inferred from ChIP seq experiments completed in and imputed for the

WA-07 female embryonic stem cell line (Ernst and Kellis 2012; Ernst and Kellis 2017). Roughly 500,000 loci fell in annotated heterochromatin regions; 22,071 and 2522 fell in weak and strong enhancers, respectively, and 2748 fell in promoters; the rest of the loci fell in other predicted chromatin states.

We hypothesized that STRs intersecting gene regulatory elements may have significantly different purity than those falling in heterochromatin regions; we assume that the latter STRs are evolving more neutrally than the former given their inferred lack of accessibility in the germline. To test this hypothesis, we first modeled the Levenshtein distance of the major allele at an STR locus as a function of whether it intersected an enhancer or heterochromatic region, after accounting for covariance in length, repeat unit size, and GC content. STRs intersecting enhancers have significantly fewer interruptions than those intersecting heterochromatic regions; furthermore, STRs are significantly purer if they intersect a strong rather than a weak enhancer (Poisson GLMs, *p* < 2 * 10^−16^, *p* = 1.46 * 10^−3^, respectively; *B* = -0.0514, -0.055, respectively) (**Figure 2A, 2B**). STRs in enhancers are also more likely to be completely pure than those falling in heterochromatin (logistic regression, *p* < 2 * 10^−16^; *B* = 0.118). We then segregated the STRs in enhancers by whether they intersected open chromatin identified by ATAC seq in iPSCs as a proxy for germline activity. STRs in enhancers that also intersected ATAC seq peaks were significantly purer than those in enhancers that did not intersect the peaks (*p* = 0.0216; B = -0.0272) (Figure 2c). We did not observe the same effect of enhancer STRs intersecting Fiber-seq Inferred Regulatory Elements (FIRE) peaks called in both GM12878 and K562 cells (*p* = 0.735) (Vollger et al. 2024).

**Figure 2.**
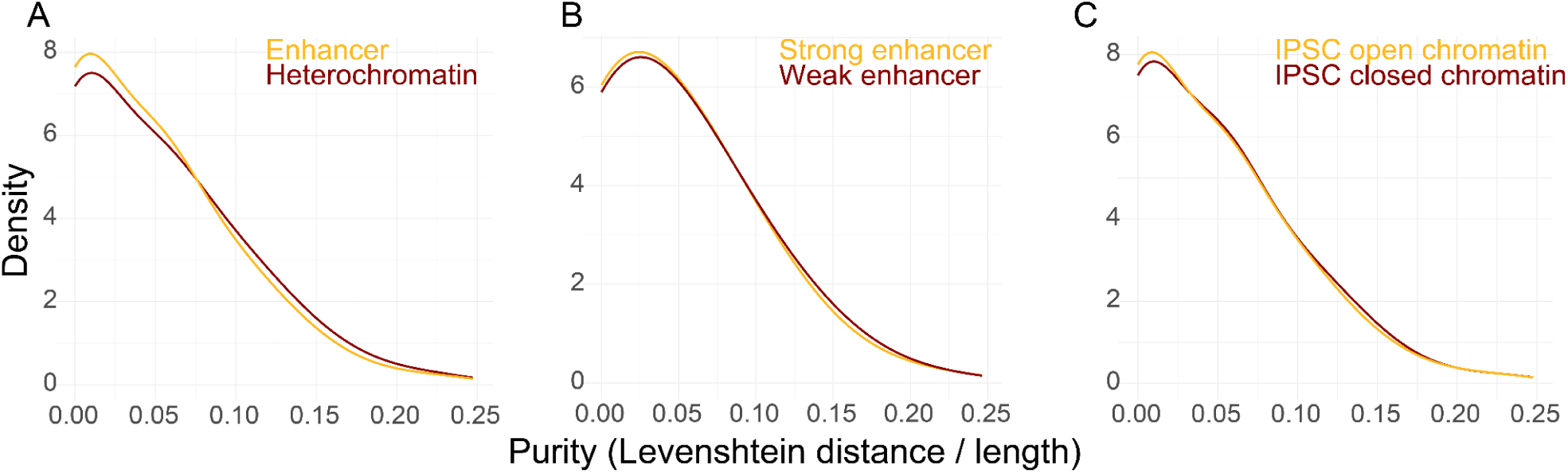
Distribution of interruptions at noncoding STRs stratified by chromatin state and regulatory function. **A**: STRs that intersect enhancers are purer than those intersecting heterochromatin. **B**: Inferred enhancer strength in ESCs is predictive of STR purity. **C**: Open chromatin in IPSCs, approximated by ATAC-seq peaks, predicts significantly purer enhancer STRs.

Interestingly, STRs that intersect promoters have even fewer interruptions than those that intersect enhancers (Poisson GLM, *p* = 0.0218, *B* = -0.038) (**Supplemental Figure 3**).

**Supplemental Figure 3:**
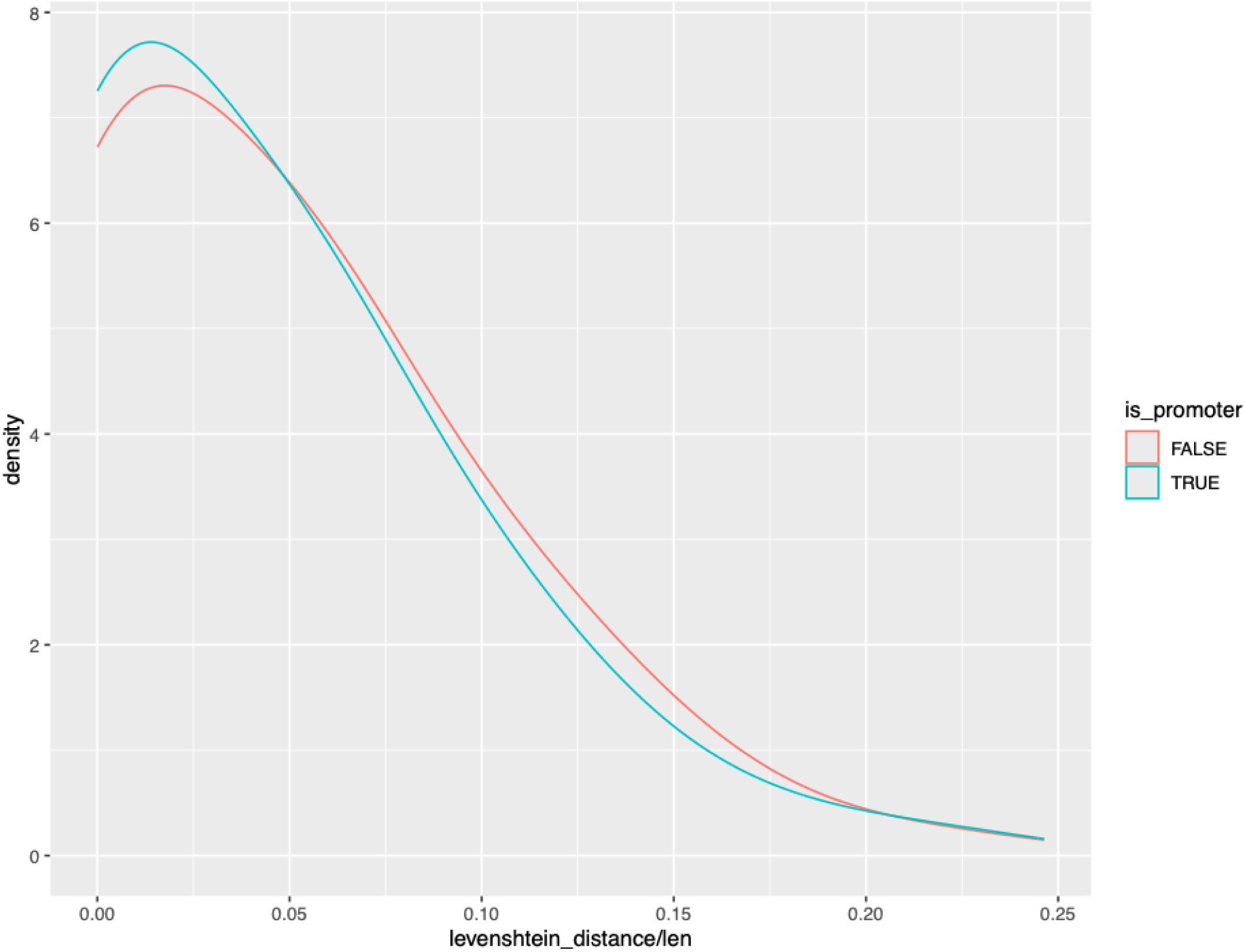
STRs that fall in promoters are significantly purer than those in enhancers.

STR mutation rate scales with the longest stretch of pure repeat; interruptions in the middle of a repeat will decrease the probability of slippage more than an interruption closer to the flanks (Wright et al. 2019; Wright et al. 2020). If interrupting an STR in a gene regulatory element incurs a roughly constant cost to protein-DNA binding activity, therefore incurring a set selective cost, we theorized that interruptions would need to maximally lower deleterious slippage to outweigh that cost. Thus, we hypothesized that, conditioning on allele purity, STRs in gene regulatory elements would maintain a shorter pure stretch of repeat. To test this hypothesis, we modeled the longest stretch of pure repeat (in bp) as a function of chromatin state while accounting for covariance with repeat unit length and GC content; as above, we used a Poisson GLM with a log link function with the log-transformed total length of the STR as an offset variable. STRs maintained a significantly longer stretch of pure repeat when intersecting an enhancer relative to intersecting heterochromatin (p < 2*10^−16^, B = 0.0141). However, when we conditioned on allele purity by including Levenshtein distance as an independent variable, we found that STRs in heterochromatin maintained a longer pure repeat (*p* = 9.65 * 10^−13^; *B* = 0.0118). AIC values indicated that the model accounting for Levenshtein distance significantly fit our data better than the model without (Δ AIC = 1034470).

STR motifs can modify their effect on DNA binding protein occupancy at neighboring binding motifs (Horton et al. 2023). Certain motifs promote significantly stronger and longer binding while others are antagonistic. In vitro experiments have mapped motif preferences for the human DNA binding protein MAX; this mapping is unknown for most other DNA binding proteins (Horton et al. 2023). AG dinucleotide repeats preferentially bind MAX while AC disfavors binding. Given our results above on purity in STRs within enhancers, we hypothesized that, with dinucleotide repeats proximal to a MAX binding site, AG STRs would be less interrupted than AC STRs. Neighboring AG STRs may be more functionally relevant to the occupancy of a MAX binding site and thus under more purifying selection. However, we observe the opposite: AG dinucleotide STRs are significantly more interrupted than AC STRs, conditional on those STRs falling in chromHMM enhancers (Poisson GLM, *p* = 0.0034, *B* = 0.547). We expanded this analysis to include nine other DNA binding proteins, including MYC, which shares a similar basic helix-loop-helix structure with MAX. Though we do not know which specific motifs promote binding for each of these proteins, we hypothesized that AC and AG dinucleotides would at minimum contrast in their effects modifying binding. After Bonferroni correction for multiple testing, interruptions varied as a function of nucleotide content for only STRs proximal to ELK1 binding sites; as above AG dinucleotide STRs were significantly more interrupted (Poisson GLM; *B* = 0.9178, Bonferroni *p* = 0.0096) (**Supplemental Figure 4**). AG dinucleotides are broadly more interrupted than AC dinucleotides at STRs near binding sites of a diverse set of DNA binding proteins; perhaps the mutational process that generates new interruptions favors AG dinucleotide repeats.

**Supplemental Figure 4:**
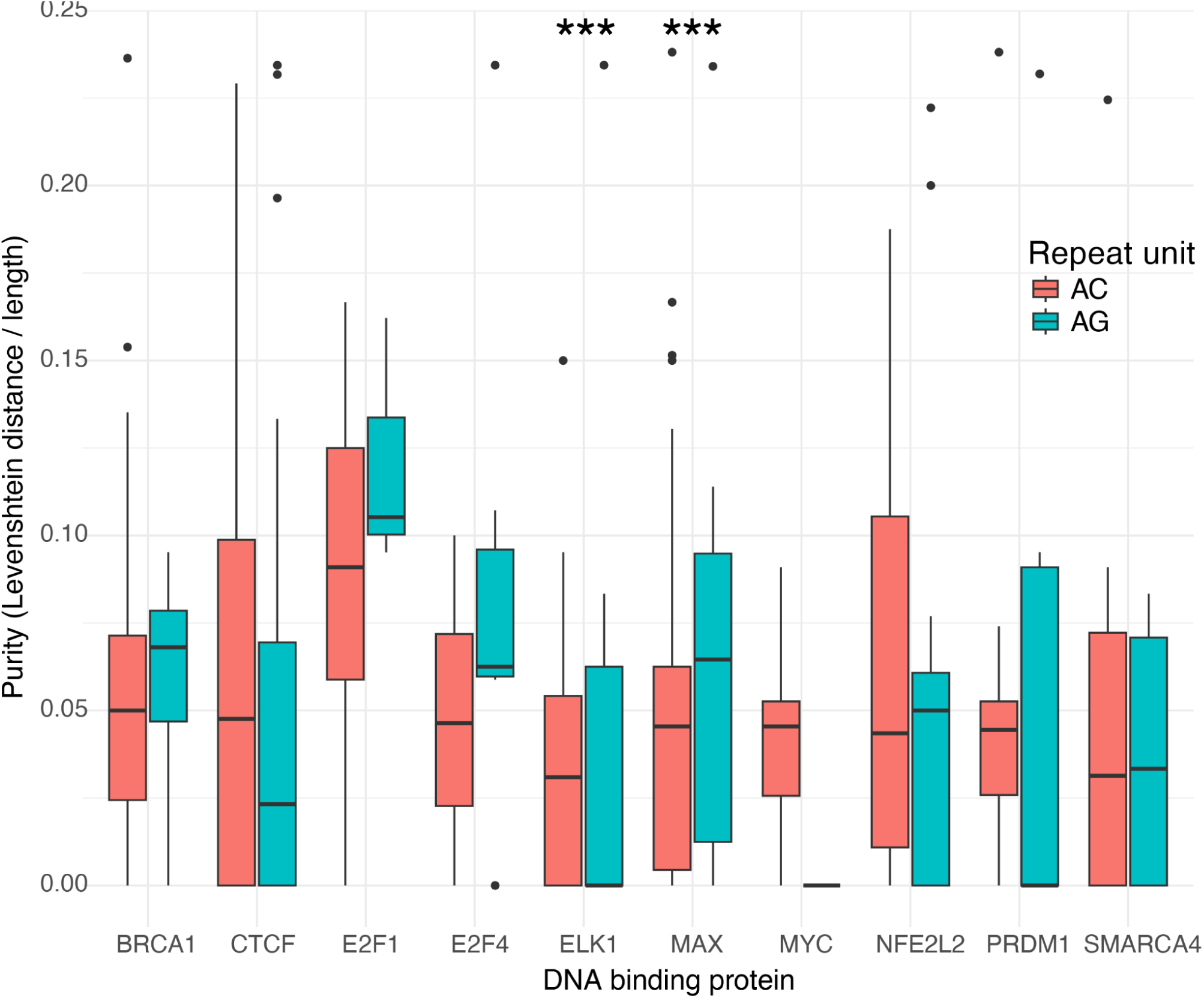
interruptions at STRs in chromHMM enhancers proximal to DNA binding protein binding sites. Asterisks indicate that AG dinucleotide repeats were significantly more interrupted than AC repeats (Bonferroni *p* < 0.05).

We next examined interruptions at enhancers identified by GeneHancer. GeneHancer integrates annotated enhancers from ENCODE, Ensembl, FANTOM, and VISTA and further infers genes whose expression they modify (Fishilevich et al. 2017).

Furthermore, the GeneHancer enhancers and their connections are rated as either “elite” or “non-elite” based on reaching a threshold of supporting evidence from multiple different experimental assays or computational methods. As before, we used Poisson GLMs with a log link function; in each model, unless otherwise specified, we controlled for the effects of allele length (as an offset variable), repeat unit length, and GC fraction (both as covariates). STRs in GeneHancer enhancers have significantly fewer interruptions than those outside (Poisson GLM, *B* = -0.035, *p* < 2 * 10^−16^). However,

STRs in GeneHancer enhancers are also significantly less likely to be pure (binomial GLM, controlling for allele length, repeat unit length, and GC fraction; *B* = -0.049; *p* = 1.12 * 10^−10^). In essence, STRs in GeneHancer enhancers are more likely to be slightly interrupted, while STRs outside of these elements are either pure or highly impure (**Supplemental Figure 5**). This nonmonotonic effect contrasts with our results using chromHMM annotations to infer gene regulatory elements. More in line with the chromHMM results, however, STRs that fall within elite GeneHancer enhancers have significantly fewer interruptions than those falling in non-elite elements (Poisson GLM, *B* = -0.017; *p* = 7.02 * 10^−4^).

**Supplemental Figure 5:**
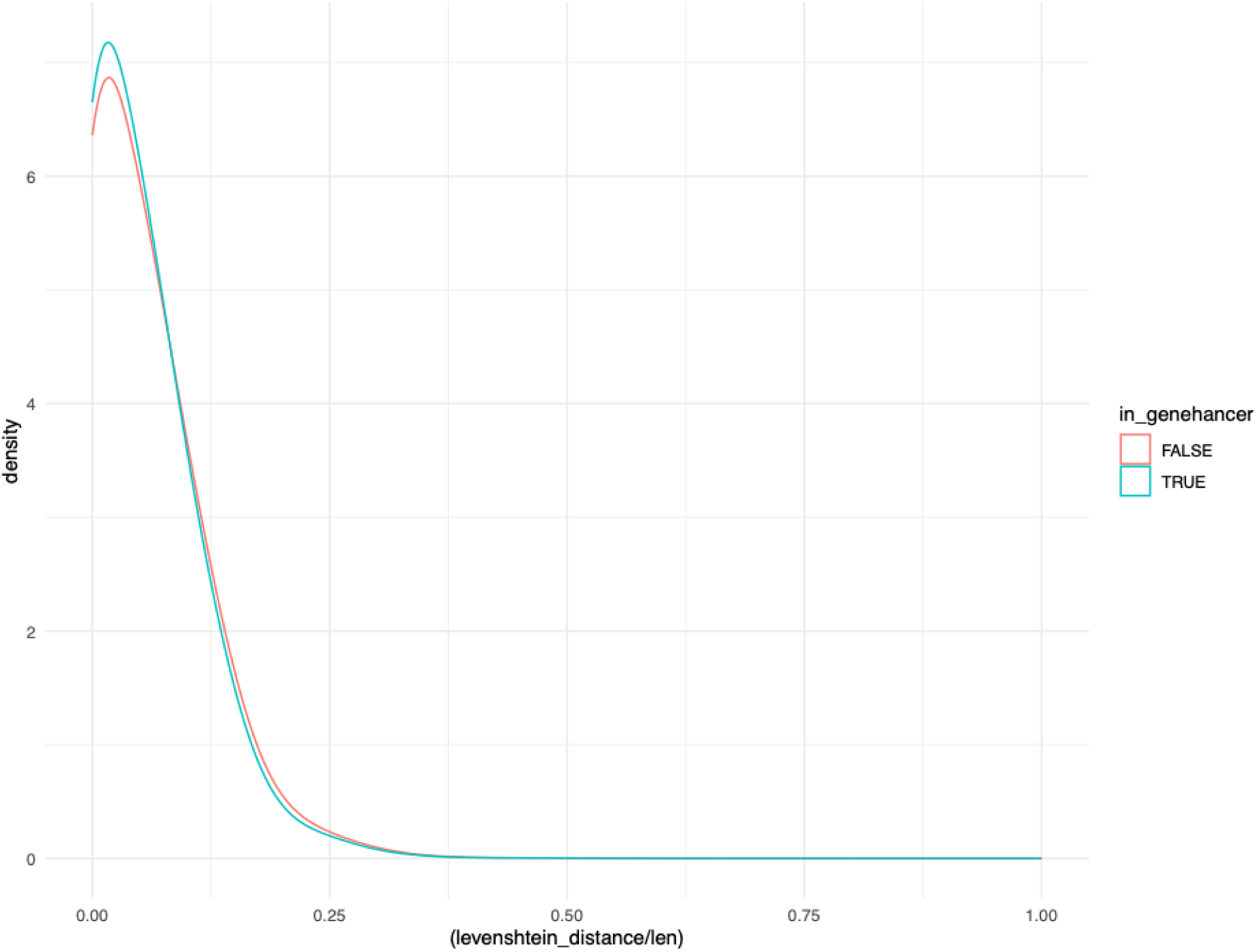
STRs in GeneHancer elements are significantly less likely to be interrupted.

GeneHancer enhancers are annotated with connections to genes whose expression they are inferred to regulate (Fishilevich et al. 2017). We leveraged these annotated gene connections to further stratify the constraint on noncoding STRs. We hypothesized that a regulatory element’s constraint could be approximated by the maximum constraint of the genes it regulates. Thus, for all STRs that overlapped elite

GeneHancer enhancers, we calculated the minimum LOEUF score of all genes with elite connections to their containing element. Higher maximum genic constraint (i.e., lower minimum LOEUF) is significantly associated with fewer interruptions at STRs that overlap elite GeneHancer enhancers (Poisson GLM, *B* = 0.032, *p* = 1.06 * 10^−4^). However, containing an elite connection to a known autosomal dominant gene did not significantly affect STR purity (Poisson GLM, *p* = 0.543).

We further examined the effect of constraint on noncoding STR interruptions using a locus-specific score of selective constraint. As described in (Mitra et al. 2021), SISTR is a method that infers the selective cost (*s*) of the gain or loss of a single motif relative to the fitness of an STR locus’s major allele. Loci where all possible allele lengths are equally fit experience unconstrained slippage and therefore high variance in allele length (*s* = 0). However, constrained loci at which slippage is deleterious have a lower variance in allele length than would be expected under neutrality (*s* > 0), as estimated by a depletion in heterozygosity observed in a population of 3200 unrelated individuals (Mitra et al. 2021). We examined 62,729 2-, 3-, and 4-mer noncoding STR loci with reliable *s* scores (**Methods**); of these, 38,146 were inferred to be under purifying selection. To infer interruptions, we counted the number of non-singleton substitution single nucleotide variants (SNVs) identified in gnomAD that fell within the boundaries of each repeat (Chen et al. 2022). After correcting for covariance with reference allele length (as an offset), GC content, and repeat unit length (both as covariates), we observed that loci under purifying selection harbored significantly fewer interrupting SNVs than those evolving neutrally (Poisson GLM with log link function, *B* = -0.0348, *p* = 4.22 * 10^−10^) (**Supplemental Figure 6**). These results appear concordant with our findings using regulatory elements to approximate constraint: higher constraint at noncoding STRs typically predicts higher purity. A caveat for this analysis, however, is that the models of neutral STR variation may not fully account for variance in purity at loci; therefore, the strength and significance of this association may be underestimated. Loci with more cryptic interruptions may harbor lower heterozygosity than expected under SISTR’s model, thus appearing under selection. As we observe the opposite association of constraint and interruptions, we are confident in the signal’s direction but the true association may be stronger than we infer.

**Supplemental Figure 6:**
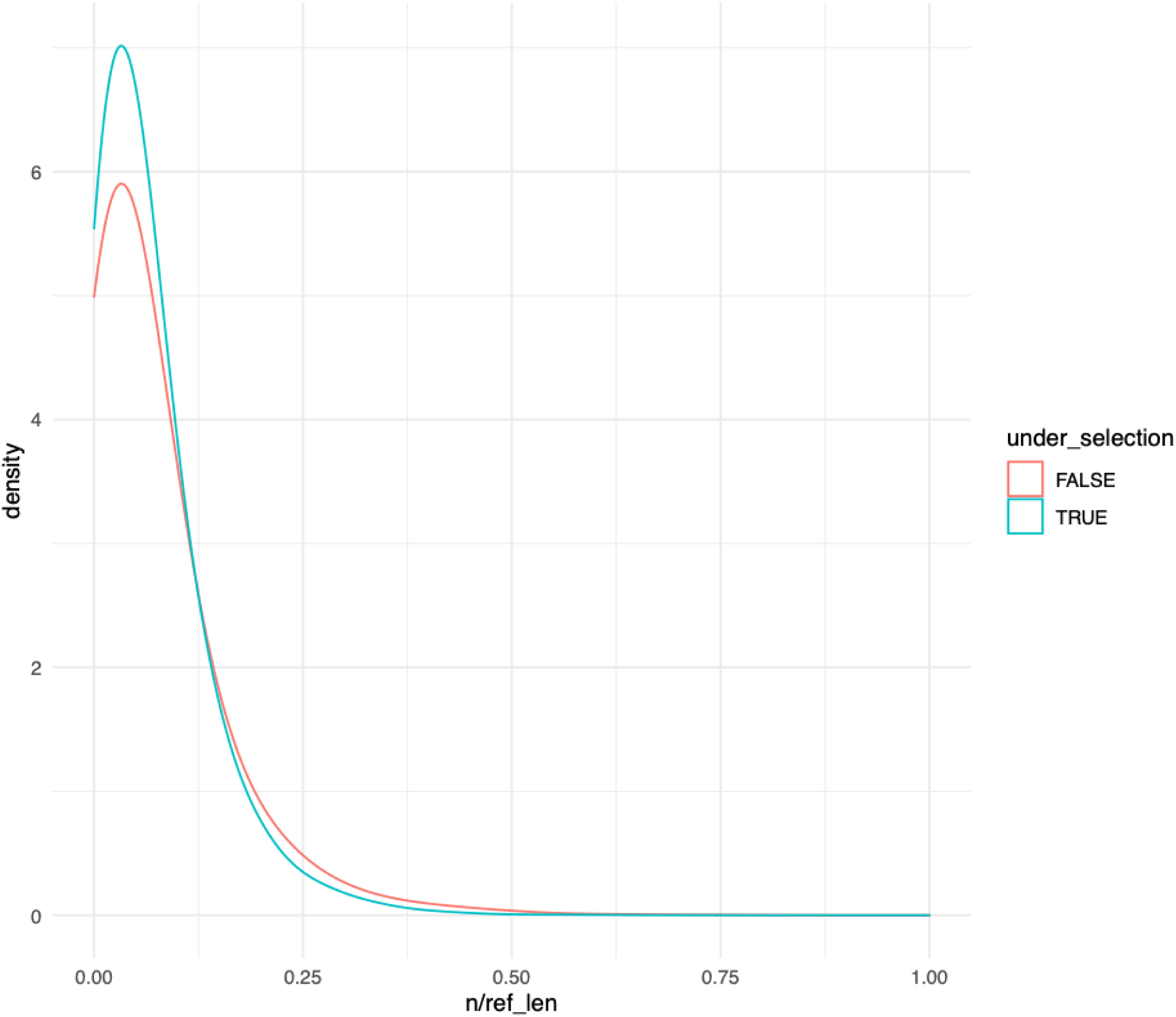
noncoding STRs under purifying selection have fewer interrupting SNVs than those evolving neutrally.

## Discussion

Allele purity is one of many characteristics that determine an STR’s instability and mutation rate (Wright et al. 2020; McGinty and Sunyaev 2023 Mar 13; Rajan-Babu et al. 2024). At loci under purifying selection, instability can lead to deleterious expanded or contracted mutant alleles; stabilizing a repeat by introducing an interruption is therefore protective against developing those deleterious alleles, as observed at a variety of loci at which expansion causes Mendelian disease (Eichler et al. 1995; Eichler and Nelson 1996; Pearson et al. 1998; Wright et al. 2019; Wright et al. 2020; Rajan-Babu et al. 2024). However, at other loci, purity may itself be under purifying selection; thus, the benefits of an interruption decreasing instability are theoretically counterbalanced by the cost of decreasing purity. At coding STRs, we observed evidence of this tradeoff affecting locus purity: at STRs in highly constrained genes, silent interruptions were favored while missense interruptions were depleted. The opposite was true at STRs in genes evolving more neutrally, at which slippage may be less deleterious and therefore increasing stability less critical. At noncoding STRs, we observe a similar tradeoff occurring. STRs in gene regulatory elements are significantly purer than those falling in putatively neutrally evolving genomic regions: in general, the cost of instability appears less than the cost of impurity.

Historically, STRs served largely as neutral markers of ancestry, encoding fine scale demographic information thanks to their high heterozygosity. More recently, however, some STRs are believed to have functional roles in phenotype. Instability at over 60 coding STR loci has been linked to Mendelian diseases; some loci in their larger noncoding counterpart are associated with complex traits and inferred to be under purifying selection (Fotsing et al. 2019; Mitra et al. 2021; Margoliash et al. 2023; Hiatt et al. 2025; Tanudisastro et al. 2025 Apr 9). The results here somewhat support this trend in recognizing that indirect positive selection may act over long periods of time to help shape locus sequence. The effects we report are significant but weak; though selection may play a role in shaping purity, mutation clearly dominates, as evidenced by the consistent strong associations of purity with allele length, repeat unit, and GC content. Future analyses may be better powered to detect effects of selection after more precisely accounting for mutation rate variation. These models could build in predicted non-B DNA structures, non-linear effects of certain motifs, or myriad other local determinants of the instability that leads to interruptions. For example, several recent studies predict nonlinear effects of allele length on instability, which theoretically implies a length-dependence in the selective benefit of an interruption (Ananda et al. 2013; McGinty et al. 2025). Even more complex models could account for both the gain and loss of interruptions through slippage (McGinty and Sunyaev 2023 Mar 13).

Furthermore, we chose proxies of STR constraint that we believed were unlikely to affect the neutral mutational processes that create new interruptions. These assumptions of independence could be incorrect. For example, if expansions generally lead to higher rates of new interruptions at STR loci in heterochromatin than in euchromatin, our interpretation of constraint explaining an observation of greater purity at loci in gene regulatory elements would be compromised. However, at minimum, the results could point to novel annotations that predict neutral mutational processes.

The large number of local determinants of instability at STRs can serve as a feature for studying their mutation rate variation: the parameter space is very large. As we observed, selection against instability tunes purity, which is just one easily alterable sequence variable that affects mutation rate. The flexible framework developed here could be easily used to test the dynamics of selection on any other alterable *cis*-acting mutation rate modifier, such as the complexity of the flanks or proximity to other STRs. Furthermore, high quality genotypes are currently being generated for the longer and more complex variable number of tandem repeats (VNTRs); the framework is similarly flexible to examine evolution of stability at these loci as altered by their unique characteristics (Nurk et al. 2022; Liao et al. 2023; Danzi et al. 2025; Yoo et al. 2025).

## Methods

For the majority of analysis, we relied on STR genotypes generated by EnsembleTR on 3550 diverse genomes from the 1000 Genomes (1KGP) and H3Africa (H3A) cohorts (The 1000 Genomes Project Consortium 2015; Choudhury et al. 2020; Byrska-Bishop et al. 2022; Jam et al. 2023). Both cohorts contain short-read WGS aligned to GRCh38; the 1KGP samples are PCR-free. EnsembleTR integrates the output of four different TR genotyping methods; of these four, HipSTR is the only tool that infers and reports the exact sequence of an allele rather than estimating its length alone; this resolution is necessary to detect interruptions (Willems et al. 2017). Thus, we subsetted the loci to those that contained HipSTR genotypes, which are limited in length to those spanned by one end of a paired-end short read (Willems et al. 2017). We excluded related individuals from the 1KGP (https://ftp.1000genomes.ebi.ac.uk/vol1/ftp/release/20130502/). We used similar filters to those applied in Jam et al., 2023 and excluded loci missing genotypes for >25% of individuals, failed Hardy-Weinberg Equilibrium testing (*p* < 0.000001), or overlapped a known segmental duplication. Individual genotypes that did not pass quality filters were also excluded. We exclusively analyzed the major allele at each locus; allele frequencies were recalculated after excluding these individuals. Homopolymer loci were excluded.

### Coding STRs

Data processing differed slightly for the coding and the noncoding STR analyses. For an STR locus to be considered “coding”, the locus needed to be fully contained within an exon from a canonical transcript, a medically-relevant exon, or an exon inferred to be under significant purifying selection; these were aggregated from RefSeq Select and MANE (Morales et al. 2022; Goldfarb et al. 2025). To calculate the number of interruptions, we calculated the number of interrupted codons relative to a pure allele, as described in the main text (**Supplemental Figure 1**). Codons differ in their “mutational space”, i.e. the fraction of possible substitutions that lead to a silent versus a missense mutation. We estimated this fraction by counting every possible substitution from the pure sequence and binned by coding consequence, excluding nonsense mutations. Major alleles at which over 25% of bases were classified as interruptions were excluded.

We used several different annotations to approximate constraint at coding STRs. Predominantly we relied on the containing gene’s LOEUF score, which is a quantitative measure of a gene’s intolerance to dominant loss-of-function mutations (Karczewski et al. 2020). Lower LOEUF scores (LOEUF ∼= 0) indicate high constraint and a depletion of observed loss-of-function mutations. We assumed here that the selective constraint on the STR would scale with that of its containing gene, as many STR expansion diseases display dominant inheritance patterns (Hiatt et al. 2025). We also intersected STRs with inferred protein domains from Pfam and downloaded the track from the UCSC Table browser (Blum et al. 2025). We used a list of genes at which deleterious alleles behave in an autosomal dominant manner generated previously (Berg et al. 2013).

### Noncoding STRs

To determine purity at noncoding STRs, we calculated the minimum Levenshtein distance between an observed allele and a pure allele, allowing for local alignment to the pure allele to account for insertions or deletions of bases other than a pure motif. All coding STRs were excluded from these analyses. As above, we examined major alleles only and excluded loci whose major allele was over 25% interrupted.

The noncoding STR analyses involved intersecting loci with a variety of annotations, as described below:

1. chromHMM: We downloaded the GRCh38 track for 25 inferred chromHMM states based on observed and imputed experiments on the WA-07 (E002) ESC cell line from https://egg2.wustl.edu/roadmap/web_portal/ (Ernst and Kellis 2012; Ernst and Kellis 2017). State annotations were downloaded from the same website. Some STR loci intersected multiple different states; all loci were classified into a single state using the following sequential logic. Loci that overlapped chromHMM positions with states 1 or 5 - 9 were classified as transcribed; states 2 - 4 were promoters; 13 - 15 were strong enhancers; 10 - 12 and 16 - 18 were weak enhancers; 19, 20, 22, or 23 were ‘other’, and the remaining loci were classified as heterochromatin.
2. DNA binding protein CHiP-seq: we downloaded conservative IDR thresholded peaks from ENCODE for ten different DNA binding proteins assayed in HeLa cells, the cell type used in Horton et al., 2023. Accession numbers and filenames for each experiment are in **Supplemental Table 1**.
3. iPSC ATAC peaks: we downloaded ATAC footprints in WTC11 iPSC cells from ENCODE (accession ENCSR506RMU).
4. FIRE peaks: we downloaded GM12878 and K562 FIRE peaks from https://stergachislab.github.io/Fiber-seq-publication-data/ (Vollger et al. 2024).

### SISTR analysis

We further analyzed the effects of noncoding STR constraint on interruptions using a set of orthogonal methods to those described above. We used SISTR scores as a measure of STR constraint (Mitra et al. 2021). SISTR infers the selective constraint at a locus by detecting lower diversity than expected given a model of mutations (expansions and contraction) under neutrality. For each locus, the score *s* refers to the expected decrease in selection coefficient associated with a difference in length of one repeat from the major allele. Scores are estimated using approximate Bayesian computation on forward-time simulations under these models of STR diversity (Mitra et al. 2021). SISTR scores are reported exclusively for 2-, 3-, and 4-mer STRs; of these, 62,729 loci had scores with a 95% confidence interval (CI) of < 0.3; higher CIs indicate a noisy estimate. Loci were classified as evolving neutrally or under constraint if their median *s* was 0 or greater than 0, respectively. Loci that overlapped coding regions were excluded.

To calculate purity, we used segregating SNVs identified in gnomAD v3 that overlapped the reference allele of the above noncoding loci (Chen et al. 2022). SNVs that did not pass quality filters or were singletons were excluded.

### Regression framework

All statistical modeling was completed in R version 4.4.2. Unless otherwise noted, regressions were Poisson GLMs with a log link function and included motif GC content and length (in bp) as covariates and allele length as an offset variable.

We used a slightly more complex model to account for codon-specific differences in mutational space while analyzing the effects of constraint on coding STRs. For any given codon, the probability that a given single base substitution would result in a silent mutation is lower than that of a missense mutation. Therefore, identifying more missense than silent interruptions could be biased by this mutational space. The number of missense interruptions for a coding allele were reweighted by the ratio of possible silent to missense single base substitutions. In Poisson GLMs that jointly modeled the distribution of both silent and missense interruptions, the offset variable for missense interruptions in a given allele was the product of that ratio and the number of codons.

## Code availability

All code to run these analyses and generate manuscript figures is available at https://github.com/goldmich/str_interruptions.

